# Iodoacetic acid disrupts mouse oocyte maturation by inducing oxidative stress and spindle abnormalities

**DOI:** 10.1101/2020.09.02.279604

**Authors:** Xiaofei Jiao, Andressa Gonsioroski, Jodi A Flaws, Huanyu Qiao

**Author notes:** Correspondence: Department of Comparative Biosciences, University of Illinois at Urbana-Champaign, 2001 South Lincoln Avenue, Urbana, IL 61802, USA.

## Abstract

Disinfection by-products (DBPs) are compounds produced during the water disinfection process. Iodoacetic acid (IAA) is one of the unregulated DBPs in drinking water, with potent cytotoxicity and genotoxicity in animals. However, whether IAA has toxic effects on oocyte maturation remains unclear. Here, we show that IAA exposure resulted in metaphase I (MI) arrest and polar-body-extrusion failure in mouse oocytes, indicating that IAA had adverse effects on mouse oocyte maturation *in vitro*. Particularly, IAA treatment caused abnormal spindle assembly and chromosome misalignment. Previous studies reported that IAA is a known inducer of oxidative stress in non-germline cells. Correspondingly, we found that IAA exposure increased the reactive oxygen species (ROS) levels in oocytes in a dose-dependent manner, indicating IAA exposure could induce oxidative stress in oocytes. Simultaneously, DNA damage was also elevated in the nuclei of these IAA-exposed mouse oocytes, evidenced by increased γ-H2AX focus number. In addition, the un-arrested oocytes entered metaphase II (MII) with severe defects in spindle morphologies and chromosome alignment after 14-hour IAA treatment. An antioxidant, N-acetyl-L-cysteine (NAC), reduced the elevated ROS level and restored the meiotic maturation in the IAA. exposed oocytes, which indicates that IAA-induced maturation failure in oocytes was mainly mediated by oxidative stress. Collectively, our results indicate that IAA exposure interfered with mouse oocyte maturation by elevating ROS levels, disrupting spindle assembly, inducing DNA damage, and causing MI arrest.

## 1. Introduction

Water disinfection byproducts (DBPs) are types of by-products produced during water disinfection processes that are designed to prevent the outbreak of waterborne diseases. DBPs can be generated from the reaction of disinfectants with organic or inorganic materials including bromide, chloride, and iodide in raw water (Fabbricino and Korshin, 2009; Farré et al., 2013). Currently, more than 700 DBPs have been detected in drinking water. Humans and non-human animals can be exposed to DBPs through various routes including inhalation, ingestion, and dermal absorption (Gonsioroski et al., 2020b; Teuschler et al., 2004). Previous studies reported that DBPs can exert adverse effects, such as cytotoxicity, genotoxicity, mutagenicity, and carcinogenicity (Richardson et al., 2007). Epidemiological studies indicate that DBP exposure is associated with the increased risk of cancers, including bladder cancer (Villanueva et al., 2004), colorectal cancer (Rahman et al., 2010), and rectal cancer (Jones et al., 2019). In addition, DBP exposure is thought to cause reproductive dysfunction (Gonsioroski et al., 2020a; Jeong et al., 2016), developmental toxicity (Andrews et al., 2004; Ding et al., 2020), and adverse pregnancy outcomes (Costet et al., 2012; Levallois et al., 2012; Toledano et al., 2005; Wright et al., 2017). The United States Environmental Protection Agency (U.S. EPA) has regulated the levels of 11 DBPs in drinking water due to the rising of health concerns (USEPA 2006). However, a large number of DBPs in drinking water are unregulated, likely due to a lack of studies on their potential toxicity.

Iodoacetic acid (IAA), an unregulated DBP in drinking water, forms during chlorinated disinfection of raw water. IAA is now attracting more attention due to its ubiquity and high toxicity. Richardson et al (2008) showed that the maximum level of IAA is up to 1.7 μg /L in water samples from 23 cities in the United States and Canada. Another study detected a high maximal level of IAA (2.18 μg/L) in drinking water in China (Wei et al., 2013).

In addition to being a ubiquitous pollutant, IAA is thought to be toxic in many systems. Wei et al (2013) showed that IAA exposure reduced cell viability in NIH3T3 cells at a concentration as low as 2.5 μM, and increased DNA damage in a concentration-dependent manner. Other studies also reported that IAA induces cytotoxicity and genotoxicity in CHO cells (Plewa et al., 2010), primary human lymphocytes (Escobar-Hoyos et al., 2013), and HepG2 cells (Wang et al., 2014; Zhang et al., 2012).

The mechanism by which IAA causes toxicity is thought to involve oxidative stress. For example, two antioxidants, catalase and butylated hydroxyanisole (BHA), reduced the percentage of IAA-induced genomic DNA damage in CHO cells by 86.5% and 42%, respectively (Cemeli et al., 2006). Antioxidants also attenuated IAA-induced cytotoxicity and genotoxicity in HepG2 cells (Wang et al., 2014) and primary cerebellar granule neurons CGNs (Zhou et al., 2015).

It is also worth noting that IAA may pose risks to female reproduction. Recent studies showed that IAA exposure inhibited follicle growth through altering the expression of the genes that regulate the cell cycle, apoptosis, estrogen receptors (ERs), and ovarian steroidogenesis (Gonsioroski et al., 2020a; Jeong et al., 2016). However, whether IAA exposure affects oocyte meiotic maturation has not been explored. Thus, in the present study, we utilized an *in-vitro* mouse oocyte culture system to study the effects of IAA exposure on oocyte maturation *in vitro*. Our results showed that IAA exposure arrested mouse oocytes at the metaphase I stage, leading to polar body extrusion (PBE) failure. IAA also increased DNA damage and disrupted spindle assembly and chromosome alignment in treated oocytes. Since IAA can induce oxidative stress in non-germline cells, we tested whether IAA-induced oxidative stress disrupts oocyte maturation. We found that IAA caused oxidative stress in mouse oocytes, which was evidenced by an elevated ROS level in response to IAA. To confirm whether oxidative stress is the major cause of the oocyte arrest, we added an antioxidant, N-acetyl-L-cysteine (NAC) in the culture media. We found that NAC rescued IAA-induced maturation failure. Collectively, the data from this study provide new information on the toxicological mechanisms underlying the effects of IAA on oocyte maturation and raise concerns that IAA exposure poses risks to reproductive health.

## 2. Materials and methods

### 2.1 Chemicals

Iodoacetic acid (≥ 98%), 3-Isobutyl-1-methylxanthine (IBMX, ≥ 99%), N-acetyl-L-cysteine (NAC, ≥ 99%) and 2′,7′-dichlorofluorescin diacetate (DCFH-DA, ≥ 97%) were purchased from Sigma-Aldrich (St. Louis, MO, USA). Pregnant mare serum gonadotropin (PMSG) was obtained from ProSpec TechnoGene.

### 2.2 Animals

CD-1 mice (Charles River Laboratories, Wilmington, MA) were used in this study and housed in the Animal Care Facility at the University of Illinois Urbana-Champaign (UIUC). Mice were housed under 12 h dark/12 h light cycles at 22 ± 1°C and were provided with food and water ad libitum. Animal handling and procedures were approved by the UIUC Institutional Animal Care and Use Committee.

### 2.3 Oocyte collection and culture

Three-week-old female mice were injected with 5 IU PMSG to stimulate follicular development. The treated mice were euthanized 48 h after PMSG treatment. Cumulus□oocyte complexes (COCs) were isolated by rupturing antral ovarian follicles with sterile needles. Cumulus cells around the fully-grown oocytes were mechanically removed as previously described (Qiao et al., 2018). The isolated oocytes were collected in M2 medium (Sigma-Aldrich, St. Louis, MO, USA) containing 100 μM IBMX. After three washes in fresh M16 medium, oocytes were cultured in a drop of pre-warmed M16 medium (Sigma-Aldrich, St. Louis, MO, USA) under mineral oil at 37°C in a 5% CO_2_ incubator. After culture in maturation medium for 2 h, 8 h, 10.5 h, and 14 h, oocytes progressed to germinal vesicle breakdown (GVBD), metaphase I (MI), anaphase I (AI)/telophase I (TI), and metaphase II (MII), respectively. Then, oocytes were collected at each stage for data analysis.

### 2.4 IAA and NAC treatment

The concentration of IAA was chosen based on our previous study (Gonsioroski et al., 2020a). IAA (10 μM and 15 μM) was found to significantly decrease follicle growth *in vitro*. Thus, we speculated that oocyte maturation may also be affected by IAA treatment because follicles are composed of oocytes and granulosa cells. IAA was dissolved in dimethyl sulfoxide (DMSO) and diluted in maturation medium (M16) with final concentrations of 2 μM, 5 μM, and 10 μM. The final concentration of the solvent DMSO was no more than 0.1% in all culture media. Oocytes in the control group were cultured in M16 with the same amount of DMSO, but no IAA. An antioxidant, NAC supplementation, was used to determine whether it rescues the damage induced by IAA. The concentration of NAC was based on a previous study by Lai et al. (2015). NAC was dissolved in M16 medium and diluted into the maturation medium with or without IAA at a final concentration of 200 μM.

### 2.5 Immunofluorescent staining

Oocytes were fixed in PBS (pH 7.4) containing 4% paraformaldehyde for 30 min at room temperature. After the oocytes were washed in washing buffer (0.1% tween-20 and 0.01% triton X-100 in PBS) twice, oocytes were permeabilized in 0.5% triton X-100 in PBS for 20 min at room temperature. After two washes, the oocytes were blocked in washing buffer with 3% BSA for 1 h at room temperature and then incubated overnight at 4°C with FITC-labelled mouse anti-α-tubulin (1:200, Sigma-Aldrich) or mouse monoclonal anti-γ-H2AX antibody, clone JBW301 (1:200, Millipore-Sigma) and anti-phospho-p44/42 MAPK (Erk1/2) (1:200, Cell Signaling Technologies). As for γ-H2AX or phospho-p44/42 MAPK (Erk1/2) staining, the oocytes were washed in washing buffer for three times (10 min each time), then incubated with the Alexa Fluor 488-conjugated goat anti-mouse IgG (H + L) and Alexa Fluor 594-conjugated goat anti-rabbit IgG (H + L) secondary antibodies (1:300, Molecular Probes) for 1 h at 37°C, respectively. Then, oocytes were washed another three times (10 min each time). Oocyte DNA was counterstained with 1 μg/ml of 4′,6-diamidine-2′-phenylindole dihydrochloride (DAPI) for 10 min at room temperature. After DAPI staining, oocytes were mounted on glass slides with 80% glycerol, examined with a Nikon A1R confocal microscope, and processed using NIS-Elements software.

### 2.6 Measuring the levels of intracellular reactive oxygen species (ROS)

To measure the ROS levels in the control and IAA-treated oocytes, 2′,7′-dichlorofluorescein diacetate (DCFH-DA) was used. Oocytes were incubated in M2 medium with 4 μM DCFH-DA for 30 minutes at 37°C in a 5% CO_2_ incubator. Then, oocytes were washed twice in PBS and the fluorescent signal was examined immediately by fluorescence microscope (Olympus, IX73). To quantify the oocyte fluorescence intensity among different treatment groups, the images of both control and IAA-treated oocytes were taken using the same imaging setup. The Image J software (National Institutes of Health, USA) was used to measure the fluorescent intensity of the regions of oocytes in these images.

### 2.7. Statistical analysis

Each treatment was performed at least three replicates. Data are presented as mean ± SEM. For comparisons between two groups, the analysis was performed using the GraphPad Prism (San Diego, USA) software followed by Student’s unpaired two-tailed t-test. Comparisons between the three groups were analyzed by one-way analysis of variance (ANOVA). Specifically, multiple comparisons between normally distributed experimental groups were performed using ANOVA followed by Tukey post-hoc comparison. Multiple comparisons between non-normally distributed experimental groups were performed using the Kruskal-Wallis test. Comparisons were considered significant at \**P* < 0.05, \*\**P* < 0.01 and \*\*\**P* < 0.001.

## 3. Results

### 3.1. Mouse oocytes failed to mature after IAA exposure

To test the effects of IAA exposure on mouse oocyte maturation, the oocytes were cultured in media supplemented with increasing concentrations of IAA (2 μM, 5 μM, and 10 μM). After 2-h or 14-h *in vitro* culture, the GVBD rate and PBE rate were recorded, respectively. As shown in Figure 1A and B, IAA exposure at all three concentrations (2 μM, 5 μM, and 10 μM) had no effect on the GVBD rate compared to control (*P* > 0.05). However, when we examined the effects of IAA exposure on PBE after 14-h culture, we found that the PBE rates of mouse oocytes were significantly reduced in the 5 μM treatment group (39.83 ± 5.54%) and 10 μM treatment group (22.11 ± 4.69%) compared to the control group (77.57 ± 6.87%)(*P* < 0.001; Fig. 1 A and C). Although the difference was not significant when compared with control, PBE decreased even when the IAA concentration was at 2 μM (70.05 ± 5.38%, *P* = 0.14). Besides, we found that this block was permanent because all of the arrested oocytes in the 14-h-culture 5-μM-IAA treatment group (Fig. 1A) still failed to extrude their polar bodies at 24 h after an additional 10-h IAA treatment, as shown in Figure 1D. Taken together, these observations revealed that IAA exposure blocked meiotic progression in mouse oocytes, which led to the failure of oocyte maturation. Since 5 μM and 10 μM IAA treatments could significantly reduce the PBE rate in mouse oocytes, these two concentrations were used for subsequent analysis to determine how IAA disrupts the oocyte maturation.

**Figure 1.**
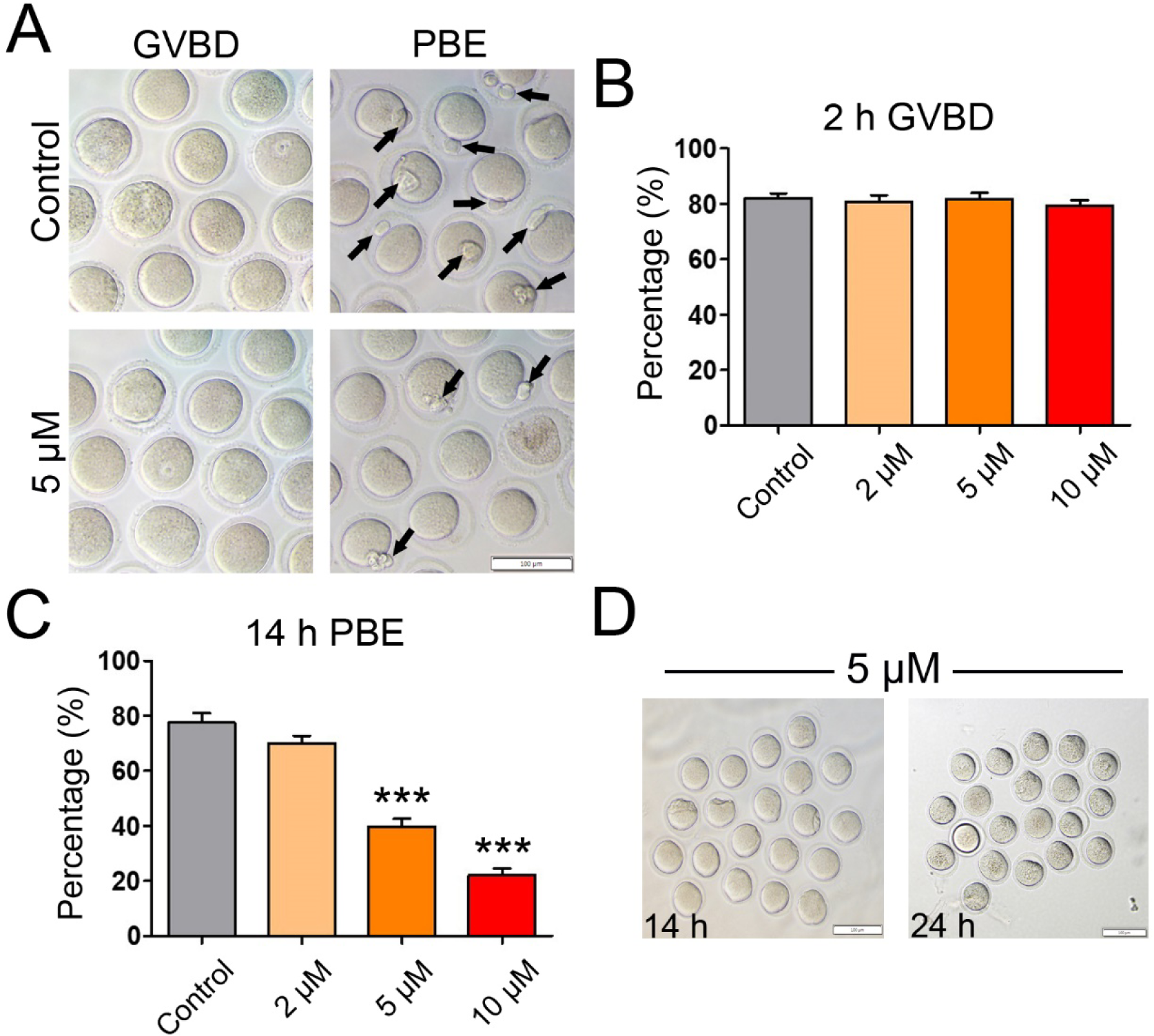
IAA exposure disrupted mouse oocyte maturation *in vitro*. (A) Representative images of oocytes with GVBD (2-h) and PBE (14-h) in the untreated control and IAA-treated groups (5 μM). The arrows highlight the oocytes with PBE. Scale bar, 100 µm. (B) The rates of GVBD in the untreated control and IAA-treatment groups (2 μM, 5 μM, and 10 μM) after 2-hour culture. (C) The PBE rates in the untreated control and IAA-treated groups (2 μM, 5 μM, and 10 μM) after 14-hour culture. (D) Representative images of the PBE-failure oocytes selected from the 5-µM-IAA-treated group that have been cultured for 14 h (left panel), continuing another 10-h culture (right panel). Note: No selected oocytes (n=20) extrude their polar bodies after 24-h culture. Scale bars, 100 µm. A total of 120 oocytes in the control, 119 oocytes in the 2 μM group, 120 oocytes in the 5 μM group, and 120 oocytes in the 10 μM group were analyzed for GVBD and PBE rate. Data were presented as mean ± SEM of at least three independent experiments. \*\*\**P* < 0.001, compared with control.

### 3.2. IAA exposure induced metaphase I arrest in mouse oocytes

Since IAA exposure had no effects on the progression of GVBD, we hypothesized that IAA exposure might result in MI arrest that finally triggered PBE failure. To determine whether the IAA induces MI arrest, we examined meiotic progression with or without IAA (5 μM and 10 μM) treatment. To stage the cultured oocytes, we immunostained the oocytes with anti-α-tubulin and counter-stained with DAPI. After 10.5-h culture (2 h for GVBD + 8.5 h), most of the cultured oocytes progressed to anaphase/telophase I (A/TI) in the untreated control group (Fig. 2A). Remarkably, the proportion of MI oocytes in the 5 μM and 10 μM IAA-treated groups was significantly higher than that in control at 10.5 h (5 μM, *P* < 0.001 and 10 μM, *P* < 0.001, Fig. 2B). On the contrary, the proportion of A/TI oocytes in IAA-treated group was significantly lower than control (5 μM, *P* < 0.001 and 10 μM, *P* < 0.001, Fig. 2B). This cell cycle analysis indicated that IAA exposure caused the oocytes to arrest at the MI stage. Further, our data also showed that IAA exposure led to MI arrest in a dose-dependent manner (Fig. 2B). In addition, this MI arrest was also verified by inverted microscopy. After 10.5-h culture, we found that over half of the control oocytes were at telophase I when polar body extrusion was processing, but only a few oocytes with forming polar bodies were found in the IAA-treated groups (Fig. 2C).

**Figure 2.**
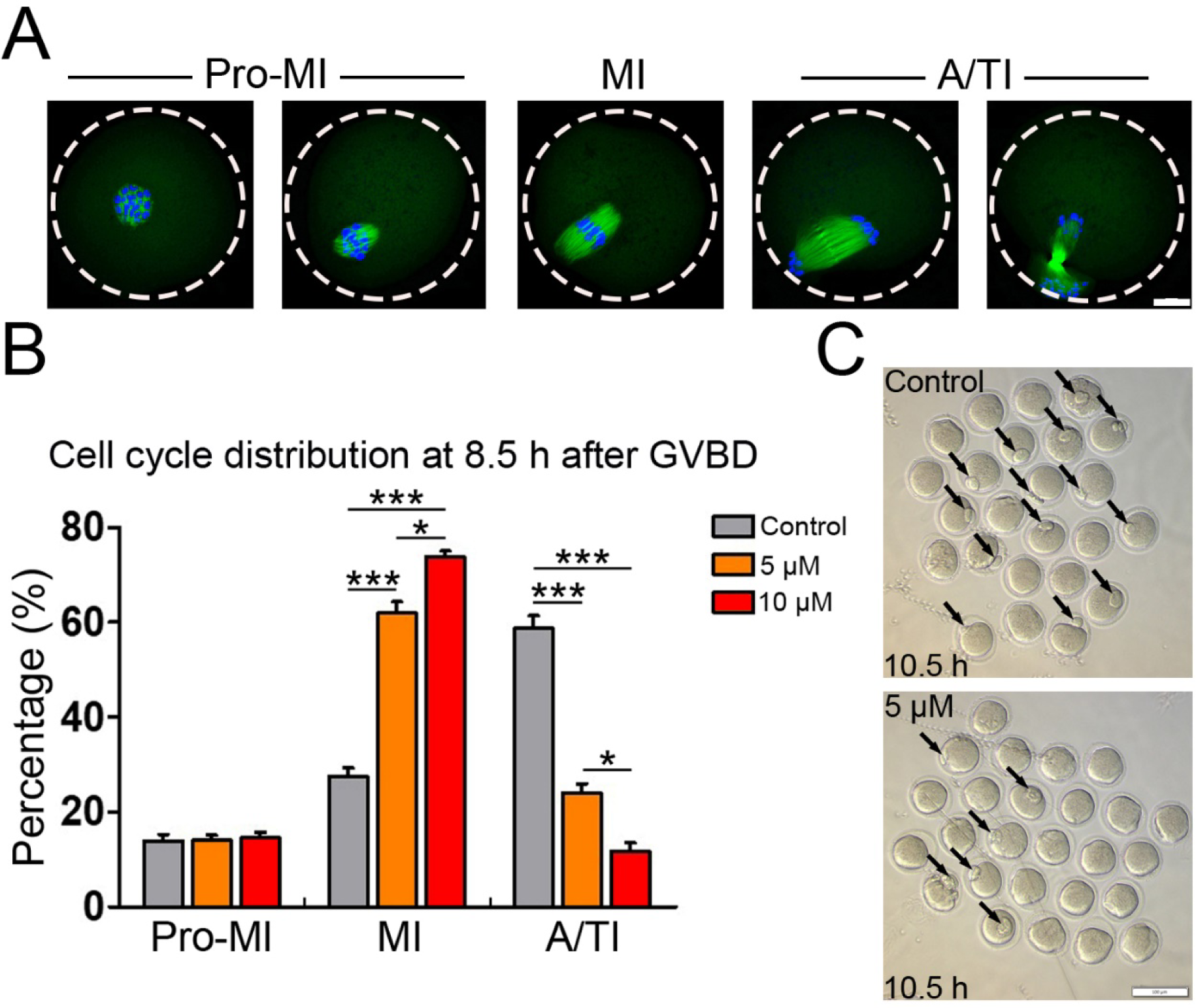
IAA exposure induced metaphase I arrest in mouse oocytes. (A) Representative images for the different meiotic stages of oocytes from pro-metaphase I (pro-MI) to anaphase/telophase I (A/TI). α-tubulin (green) was immunostained to show spindle morphology. 4′,6-diamidine-2′-phenylindole dihydrochloride (DAPI) stained chromosomes/DNA (blue). Scale bar, 20 μm. (B) The proportions of different cell stages (pro-MI, MI, A/TI) in the control, 5 μM and 10 μM IAA-treated groups. A total of 66 oocytes in the control, 71 oocytes in the 5 μM group and 69 oocytes in the 10 μM group were used for cell cycle analysis. (C) Representative images show the morphologies of oocytes in the IAA (5μM)-treated and control groups. Arrows point to the forming polar bodies. Scale bar, 100 μm. \*\*\**P* < 0.001, compared with control. \**P* < 0.05, 5 μM versus 10 μM.

### 3.3. IAA exposure caused abnormalities of spindle assembly and chromosome alignment in MI mouse oocytes

To understand how IAA induces PBE failure in oocytes, we investigated spindle assembly and chromosome alignment during MI because these processes are required for PBE. Unexpectedly, spindle assembly was not extremely disrupted by IAA (5 μM and 10 μM) exposure as evidenced by a barrel-shaped spindle (Fig. 3A). Next, we measured the spindle length and width of oocytes in control, 5 μM and 10 μM IAA-treated groups. IAA exposure significantly increased spindle length in a dose-dependent manner (36.64 ± 4.32 μm in 5 μM IAA; 39.98 ± 4.92 μm in 10 μM IAA versus 32.03 ± 3.58 μm in control, *P* < 0.001; 5 μM IAA versus 10 μM IAA, *P* < 0.01; Fig. 3B), and it significantly decreased spindle width (17.95 ± 1.82 μm in 5 μM IAA, *P* < 0.01; 17.48 ± 2.57 μm in 10 μM IAA, *P* < 0.001 versus 19.55 ± 1.91 μm in control; Fig. 3C). In addition, 5 μM and 10 μM IAA did not induce severe dispersed distribution of chromosomes as often found in arrested oocytes (Fig. 3A). However, misalignment of meiotic chromosomes was observed after we measured the MI-plate width. We found that IAA induced a significant increase in MI-plate width compared to the control (12.07 ± 2.45 μm in 5 μM IAA, *P* < 0.01; 13.59 ± 2.80 μm in 10 μM IAA, *P* < 0.001 versus 10.71 ± 1.86 μm in control; Fig. 3D). In addition, we found that a high IAA concentration (30 μM) caused severe defects of spindle formation and chromosome alignment in treated oocytes (Fig. S1). Particularly, spindles in the 30-μM-IAA-treated oocytes were extremely long and disorganized, which was evidenced by the mislocalization of p-ERK1/2, microtubule organizing centers (MTOCs) associated proteins that enrich at spindle poles to facilitate proper spindle formation (Fan and Sun, 2004).

**Figure 3.**
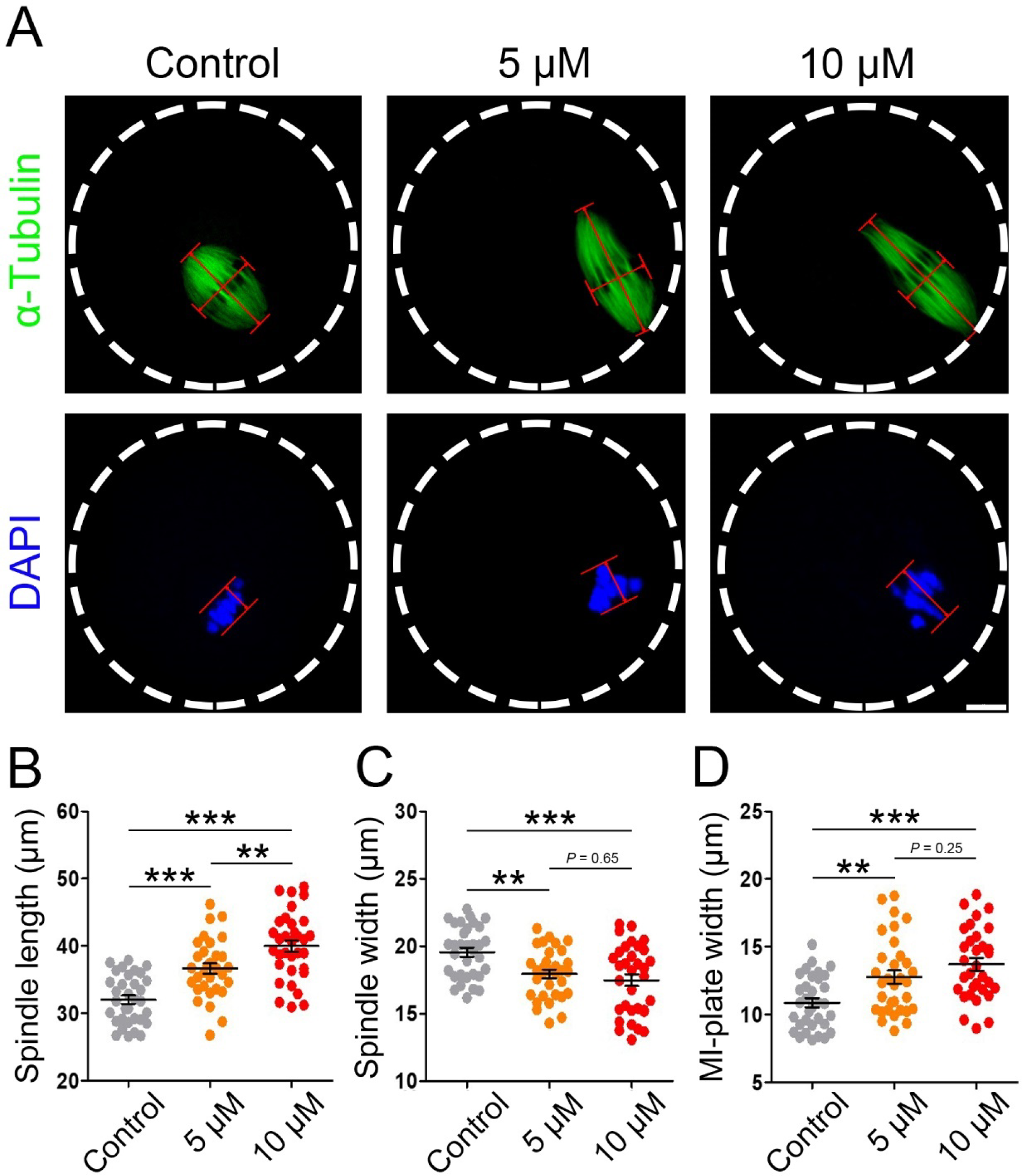
IAA exposure induced abnormalities of spindle assembly and chromosome alignment in MI mouse oocytes. (A) Representative images of MI spindle morphologies in the control, 5 μM and 10 μM IAA-exposed oocytes. Spindle length was calculated by the distance from one spindle pole to the other. Spindle width was represented by the width of microtubules at the MI plate. MI plate width was measured by the axis distance between the two lines at the edges of the DNA. α-tubulin, green; chromosomes/DNA, blue. Scale bar, 10 μm. (B) Quantitative analysis (mean ± SEM) of the meiotic spindle length in three treatment groups. (C) Quantitative analysis (mean ± SEM) of the meiotic spindle width in three treatment groups. (D) Quantitative analysis (mean ± SEM) of the MI-plate width in three treatment groups. A total of 33 oocytes in the control, 32 oocytes in the 5 μM group and 33 oocytes in the 10 μM group were used for spindle length and width analysis; A total of 33 oocytes in the control, 32 oocytes in the 5 μM group and 32 oocytes in the 10 μM group were used for MI-plate width analysis. \**P* < 0.01, \*\*\**P* < 0.001, compared with control. \**P* < 0.01, 5 μM versus 10 μM.

### 3.4. IAA exposure increased the level of reactive oxygen species (ROS) in mouse oocytes

IAA was reported to induce an elevation of ROS, leading to oxidative stress in mammalian cells (Wang et al., 2014; Zhou et al., 2015), and oxidative stress is well-known to be detrimental to oocyte maturation (Combelles et al., 2009). Thus, we hypothesized that IAA exposure results in increased ROS levels in mouse oocytes, which may lead to the aforementioned defects. To test our hypothesis, we used an oxidation-sensitive fluorescent probe, DCFH-DA, to monitor the generation of intracellular ROS in mouse oocytes. Our results showed that the DCFH-DA signals in 5 μM and 10 μM IAA-treated oocytes were brighter than in control oocytes (Fig. 4A). Our quantitative analysis verified the observation: the fluorescence intensity of DCFH-DA in 5 μM and 10 μM IAA treated oocytes was significantly higher than that in control oocytes (*P* < 0.001, Fig. 4B). Meanwhile, IAA raised intracellular ROS levels in oocytes in a dosage-dependent manner, the DCFH-DA intensity in the 10 μM IAA-treated oocytes was significantly higher than in the 5 μM group (*P* < 0.05, Fig. 4B).

**Figure 4.**
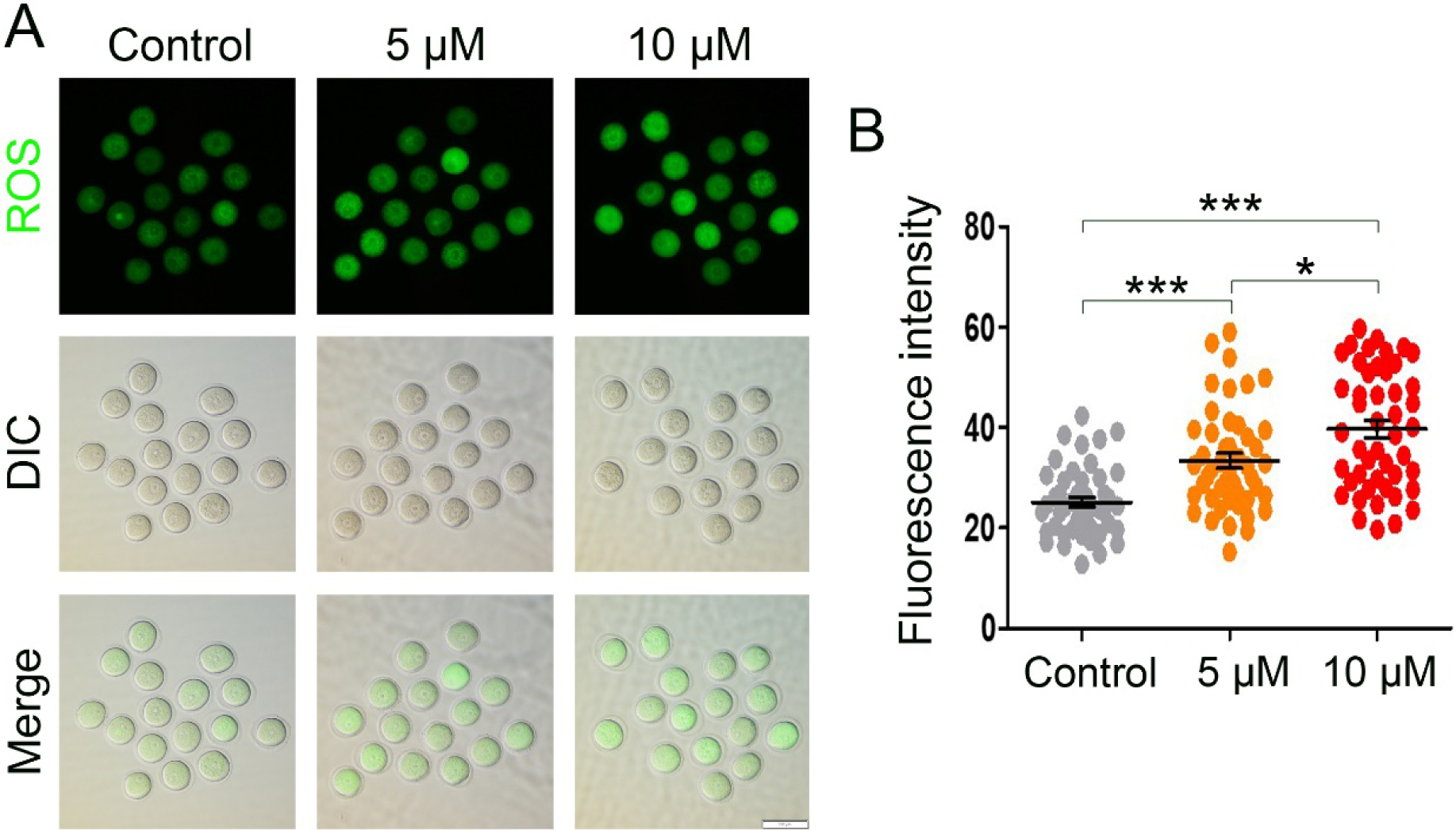
IAA exposure increased the ROS level in mouse oocytes. (A) Representative images of DCHF-DA fluorescence (green) in the control and IAA (5 μM and 10 μM)-treated oocytes. DIC, differential interference contrast. Scale bar, 100 μm. (B) Quantitative analysis of DCHF-DA fluorescence intensity in the control and IAA (5 μM and 10 μM)-exposed groups. A total of 46 oocytes in the control, 47 oocytes in the 5 μM group, and 47 oocytes in the 10 μM group were used for analysis. Data were presented as mean percentage (mean ± SEM) of at least three independent experiments. \*\*\**P* < 0.001, compared with control. \**P* < 0.05, 5 μM versus 10 μM.

### 3.5. IAA exposure induced DNA damage in mouse oocytes

ROS can drive genomic instability and elevated ROS levels usually lead to DNA damage (Salehi et al., 2018; Srinivas et al., 2019). Thus, we hypothesized that IAA exposure results in DNA damage in mouse oocytes due to an increase of cellular ROS levels after IAA exposure. We further checked the DNA double-strand breaks (DSBs) in GV-arrested oocytes using γ-H2AX staining. IBMX was used to arrest the oocytes at the GV stage for 8 h in the three treatment groups (5 μM IAA, 10 μM IAA, and the untreated control). Our results show that more γ-H2AX foci were found in 5 μM and 10 μM IAA-treated oocytes compared to the control (Fig. 5A). Next, comparison between the three groups was made using the Kruskal-Wallis test, and our results showed that the average focus number in 5 μM and 10 μM IAA-treated oocytes was significantly higher than in control (2.94 ± 1.96 in 5 μM, *P* < 0.05; 4.16 ± 2.10 in 10 μM, *P* < 0.01 versus 1.48 ± 1.50 in control) (Fig. 5B). Meanwhile, IAA induced oocyte DNA breaks in a dose-dependent manner and the average foci number in 10 μM IAA-treated oocytes was significantly higher than in the 5 μM group (*P* < 0.05, Fig. 5B).

**Figure 5.**
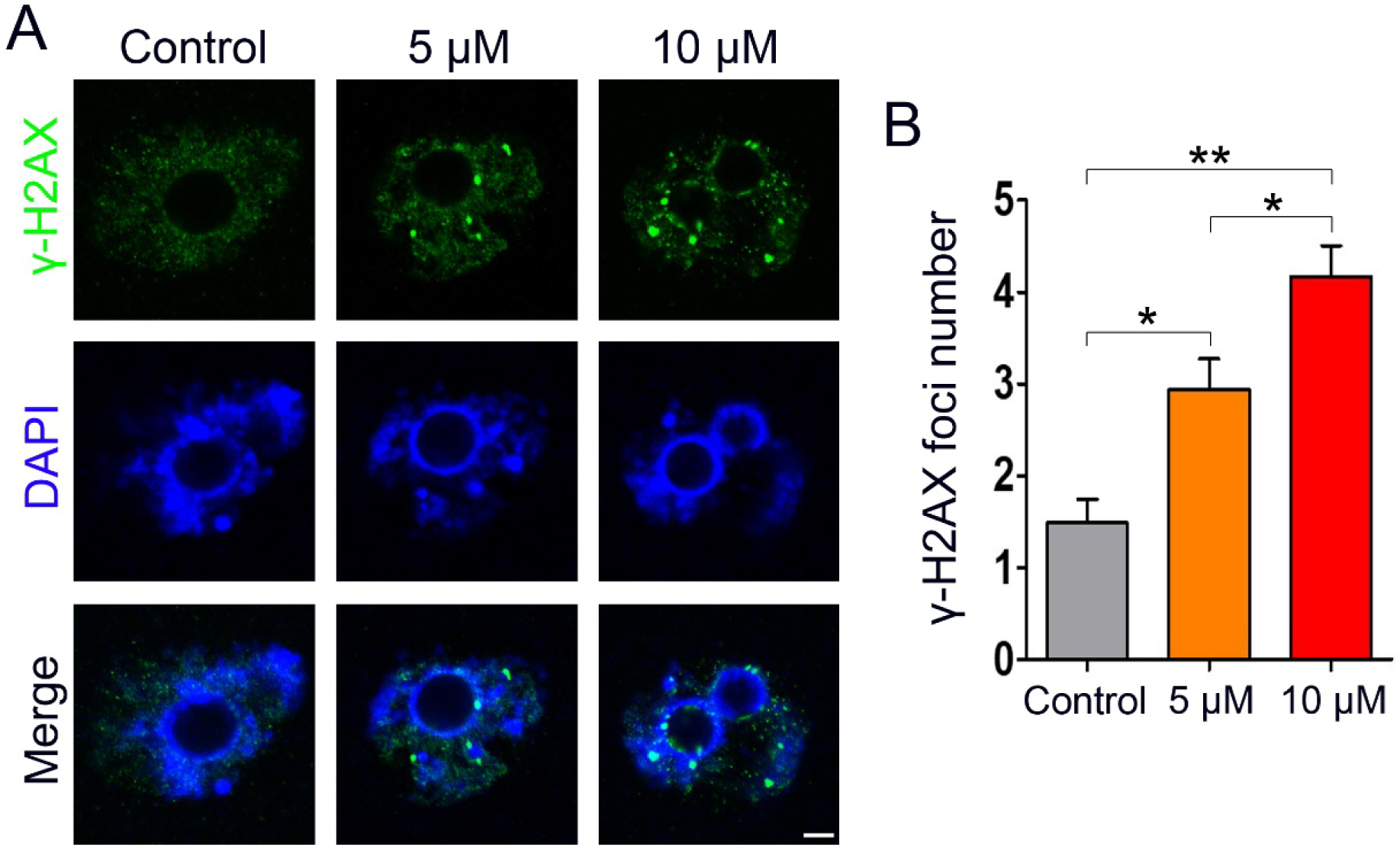
IAA exposure induced DNA damage in mouse oocytes. (A) Oocytes were arrested at the GV stage by IBMX for 8 h with or without IAA exposure, and then these GV-arrested oocytes were collected for DNA damage detection. Representative images of oocytes with DNA double-strand breaks (DSBs) indicated by γ-H2AX foci. Normal oocytes exhibited no or a few foci, whereas more foci can be found in the IAA-exposed oocytes. Green, γ-H2AX; blue, DNA. Scale bar = 5 μm. (B) Quantification of γ-H2AX foci in each group. A total of 33 oocytes in the control, 36 oocytes in the 5 μM group, and 37 oocytes in the 10 μM group were used for analysis. Data were presented as mean percentage (mean ± SEM) of at least three independent experiments. \**P* < 0.05, \*\**P* < 0.01, compared with control; \**P* < 0.05, 5 μM versus 10 μM.

### 3.6. IAA exposure impaired spindle formation and chromosome alignment in metaphase II oocytes

After IAA exposure, some oocytes could still undergo PBE. To check the quality of these surviving oocytes, we examined the spindle morphologies of MII oocytes in the 5 μM IAA-treated group. Our results showed that most of the MII oocytes in the control group displayed a typical spindle shape and well-aligned chromosomes, whereas many of the 5 μM IAA-treated MII oocytes had abnormal spindle morphologies and misaligned chromosomes (Fig. 6A). Statistically, the proportion of oocytes with abnormal spindles in the IAA-treated group was significantly higher than that in the control (39.28 ± 4.27% in 5 μM IAA versus 15.68 ± 9.25% in control, *P* < 0.01; Fig. 6B). In addition, the proportion of oocytes with misaligned chromosomes in the IAA-treated group was significantly higher than that in control (32.10 ± 6.36% in 5 μM IAA versus 13.39 ± 4.90% in control, *P* < 0.01; Fig. 6C).

**Figure 6.**
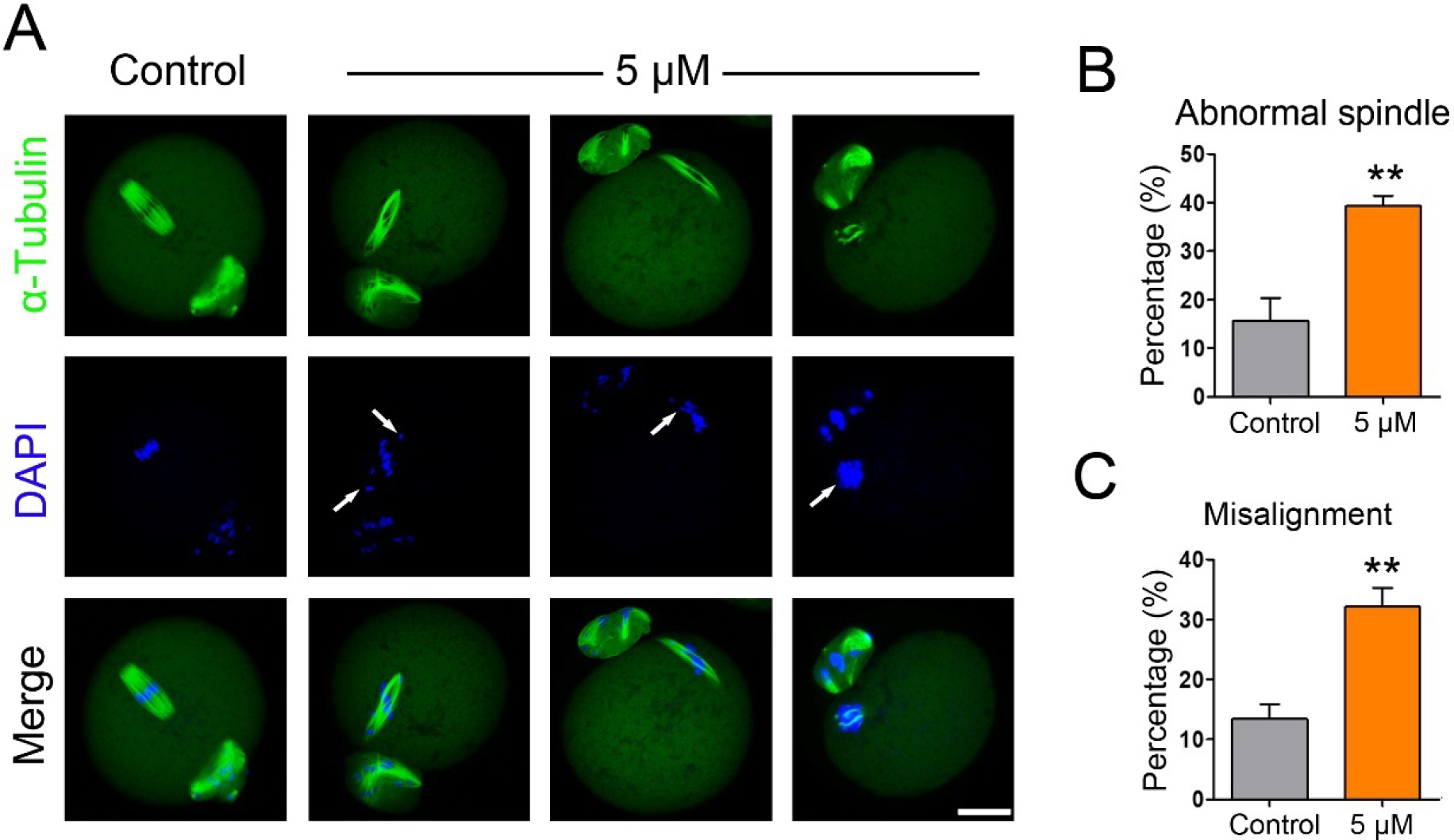
IAA exposure disrupted MII spindle formation and chromosome alignment. (A) Representative images of spindle morphology and chromosome alignment in control and IAA-treated MII oocytes. α-tubulin, green; chromosomes/DNA, blue. Scale bar, 20 μm. Arrows highlight the misaligned chromosomes. (B) The percentage of oocytes with abnormal spindles in the control and 5 μM IAA-treated groups. (C) The proportion of oocytes with misaligned chromosomes in the control and 5 μM IAA-treated groups. A total of 54 MII oocytes in the control and 54 in the 5 μM IAA-treated group were used for analysis. Data were presented as mean percentage (mean ± SEM) of at least three independent experiments. \*\**P* < 0.01, compared with control.

### 3.7. An antioxidant, N-acetyl-L-cysteine (NAC), improved the meiotic maturation rate in IAA□exposed oocytes

To investigate whether IAA-induced oxidative stress can be alleviated by an antioxidant, NAC (200 μM), we examined ROS levels in the control, 5 μM, and 10 μM IAA-treated oocytes with and without NAC treatment. We found that NAC significantly reduced the elevated ROS level in the 5 μM and 10 μM IAA-treated oocytes (5 μM, *P* < 0.01 and 10 μM, *P* < 0.001, Fig. 7A and B). Subsequently, the PBE rates were analyzed in all treatment groups to test whether the antioxidant NAC can further rescue the PBE failure induced by IAA exposure. We observed that more oocytes successfully extruded the polar bodies in NAC + IAA co-treated groups compared to the IAA-only groups (Fig. 7C). We also found the PBE rates in the 5 μM and 10 μM IAA groups were significantly lower than in the corresponding NAC + IAA groups (*P* < 0.01, Fig. 7D).

**Figure 7.**
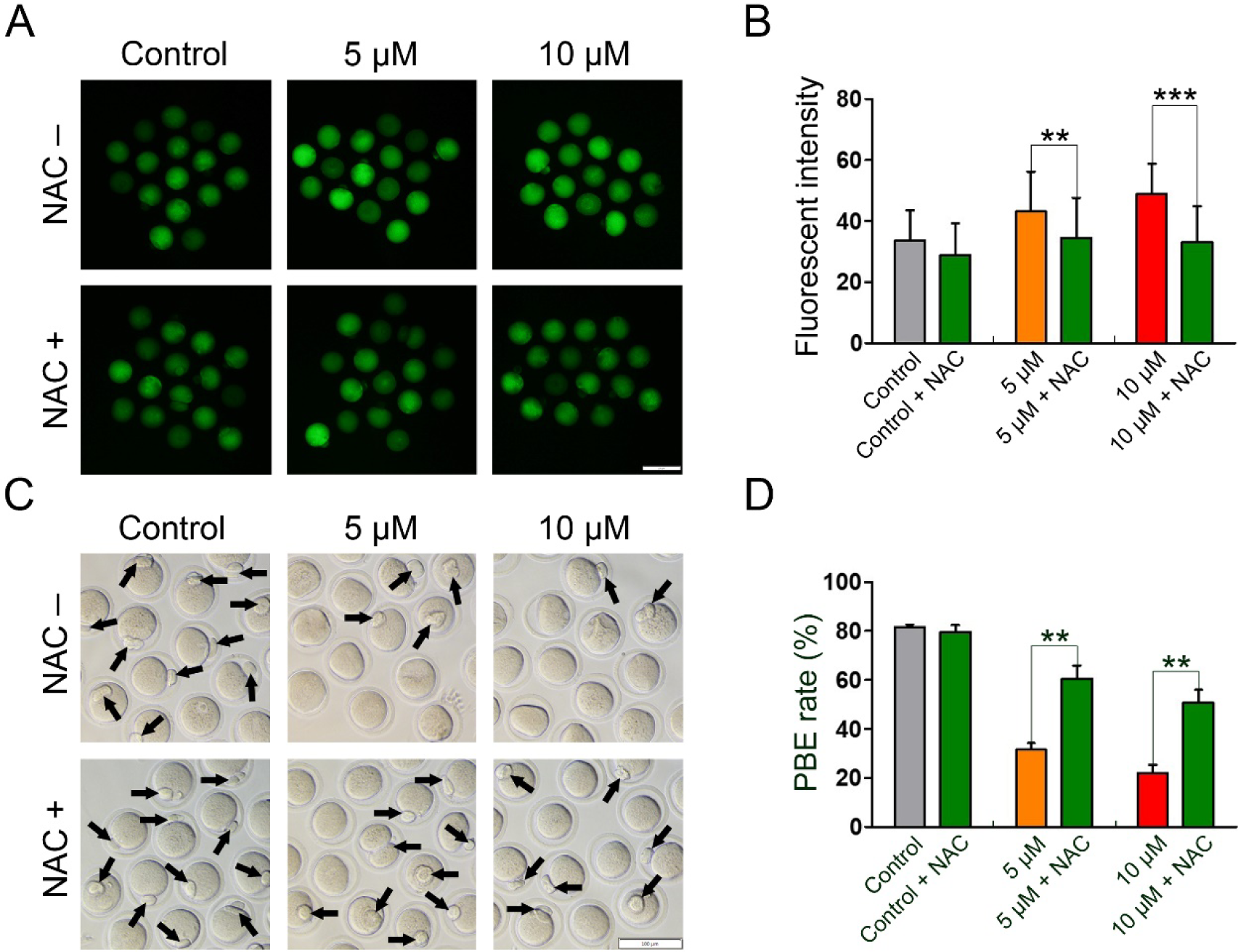
NAC rescued the meiotic arrest in IAA□exposed mouse oocytes by reducing oxidative stress. (A) Representative images of ROS levels in the control, IAA-treated, and NAC+IAA-treated oocytes. Scale bar, 100 μm. (B) Fluorescence intensity of ROS in the oocytes from different treatment groups. A total of 31 oocytes in the control, 32 oocytes in the control + NAC, 33 oocytes in the 5 μM group, 32 oocytes in the 5 μM + NAC group, 32 oocytes in the 10 μM group, and 32 oocytes in the 10 μM + NAC group were used for ROS analysis. (C) Representative images of oocytes with PBE (14-h) in the control, 5 µM, and 10 µM IAA-treated groups with and without NAC treatment. The arrows highlight the oocytes with PBE, Scale bar, 100 µm. (D) The PBE rates in the control, 5 µM, and 10 µM IAA-treated groups with and without NAC treatment after 14-hour culture. A total of 65 oocytes in the control, 59 oocytes in the control + NAC, 73 oocytes in the 5 μM group, 73 oocytes in the 5 μM + NAC group, 69 oocytes in the 10 μM group and 69 oocytes in the 10 μM + NAC group were used for PBE analysis. Data were presented as mean percentage (mean ± SEM) of at least three independent experiments. \*\**P* < 0.01, \*\*\**P* < 0.001; 5 μM versus 5 μM + NAC; 10 μM versus 10 μM + NAC.

## 4. Discussion

Recent studies showed that IAA, an unregulated water disinfection by-product, inhibited antral follicle growth *in vitro* (Gonsioroski et al., 2020a; Jeong et al., 2016), which suggests that IAA is a reproductive toxicant. This is a concern because female fertility is declining in the United States (Hamilton et al., 2006) and many environmental toxicants have been identified as exogenous factors that are associated with a decline of female fertility (Hunt and Hassold, 2008; Rattan et al., 2017; Sifakis et al., 2017). Particularly, oocytes are susceptible to chemical-induced perturbations. Oogenesis along with follicle development produces oocytes that are essential for female reproduction, but little was known about the effects of IAA on oocyte maturation. Thus, to our knowledge, this is the first study to explore IAA effects on oocyte maturation as well as the possible mechanisms underlying IAA toxicity on oocytes *in vitro*. We have documented the occurrence of PBE failure, abnormal spindle assembly, chromosome misalignment, ROS level elevation, and DNA damage in mouse oocytes in response to IAA exposure, and uncovered that the toxic effects were mainly mediated by IAA induced-oxidative stress.

GVBD and PBE are two markers of meiotic progression during oocyte *in-vitro* maturation (IVM). The first step of IVM is isolating oocytes from antral follicles. The isolated oocytes are at the germinal vesicle (GV) stage and they will resume meiosis after *in-vitro* culture. The germinal vesicle/nuclear envelope will subsequently breakdown. This GVBD process can be morphologically characterized by microscopy (Pan and Li, 2019). When oocytes progress from GVBD to metaphase I stage, the connections between spindle and chromosomes are well established. After the segregation of homologous chromosomes, the oocyte will extrude the first polar body and quickly arrest at the metaphase II stage where the oocyte can be fertilized. Therefore, the extrusion of the first polar body usually marks oocyte maturation (Sánchez and Smitz, 2012). Defects in GVBD will affect the subsequent meiotic progression, which could finally delay or block oocyte maturation (Jiao et al., 2017; Zhang et al., 2020). Another timepoint that can determine oocyte maturation is the metaphase I to anaphase I (MI-AI) transition. Abnormal meiotic events during MI would lead to MI arrest and subsequently inhibit the extrusion of the polar body. Our results showed that IAA did not affect the GVBD rate at any treatment concentrations (Fig. 1B), suggesting that IAA is not toxic to GVBD in mouse oocytes at the selected concentrations. However, IAA (5 μM and 10 μM) significantly reduced PBE rates after 14-h culture in a concentration-dependent manner (Fig. 1C), indicating that IAA is toxic to PBE in oocytes at the selected doses. Since GVBD was not disturbed by IAA, we hypothesized that IAA might arrest most of the oocytes at MI, even though a few of oocytes would proceed to the MII stage. Cell cycle analysis supported our hypothesis because more IAA-treated oocytes arrested at the MI stage compared to the untreated control (Fig. 2). These arrested oocytes in the IAA (5 μM)-treated group could not progress further to undergo PBE at 24 h. Taken together, IAA-exposure induced MI arrest that finally resulted in the failure of oocyte maturation.

Proper spindle assembly is essential for chromosome segregation, oocyte maturation, and subsequent embryo development (Howe and FitzHarris, 2013). Our results showed that IAA (5 μM and 10 µM) exposure did not cause severe disruption of spindle assembly in oocytes as evidenced by the bipolar-barrel morphology (Fig. 3), however, 5 μM and 10 μM IAA increased spindle length and decreased spindle width, indicating that IAA disrupted normal spindle morphology by altering spindle shapes (Fig. 3). In addition, IAA impaired chromosome alignment as evidenced by increased MI-plate width. We also found that 30-µM-IAA treatment resulted in very long spindles (Fig. S1), which further verified that IAA exposure elongates spindle length in oocytes. It is worth noting that balance between microtubule (MT) polymerization and depolymerization is important for spindle length control and knockdown of some MT depolymerization-related proteins will cause elongated spindles (Do et al., 2014; Wang et al., 2010). Thus, we speculated that IAA (5-30 μM in our study) might disrupt the MT depolymerization in oocytes, which further study is needed to verify this hypothesis.

It is known that spindle assembly checkpoint (SAC) is insufficient for preventing chromosome mis-segregation in mammalian oocytes (Sebestova et al., 2012). Thus, only extreme spindle damage and chromosome misalignment can induce MI arrest to prevent polar body formation. Therefore, we hypothesized that the IAA (5 μM and 10 μM)-induced mild abnormality of spindle assembly and chromosome alignment is not sufficient to trigger robust MI arrest. In addition, our data indicated that 30 μM IAA did cause severe spindle defects and chromosome misalignment (Fig. S1), which suggests that high IAA concentrations might induce MI arrest in oocytes. Additionally, we observed that the IAA-treated oocytes had more severely misshaped spindles at MII compared to MI. Therefore, we inferred that a longer IAA exposure would increase the severity of spindle mis-assembly and chromosome misalignment. Since abnormal MII spindle is associated with poor pregnancy outcome (Kilani et al., 2009), IAA also reduced the quality of matured oocytes.

Previous studies have associated IAA exposure with oxidative stress (Cemeli et al., 2006; Wang et al., 2014; Zhou et al., 2015). IAA exposure increased the ROS levels and induced Nrf2-mediated antioxidant response in HepG2 cells (Wang et al., 2014). IAA also elevated the ROS levels that cause oxidative stress in primary cerebellar granule neurons (Zhou et al., 2015) and CHO cells (Cemeli et al., 2006). Consistent with these previous studies, our data show that IAA increased ROS levels in mouse oocytes in a dose-dependent manner (Fig. 4). Since excessive ROS generation will result in oxidative stress that can impede oocyte maturation (Jiao et al., 2019, 2020), we hypothesized that IAA-induced ROS elevation inhibits oocyte maturation *in vitro*. Excessive ROS generation is also associated with causing DNA damage in cells (Salehi et al., 2018; Srinivas et al., 2019). Our results show that IAA treatment induced a significantly higher level of DNA damage compared to controls as indicated by γ-H2AX focus formation, a marker for DNA double-strand breaks (Fig. 5). This result is consistent with other IAA-toxicological studies, such as research in CHO cells (Plewa et al., 2010), in human peripheral blood lymphocytes and sperm (Ali et al., 2014), as well as in NIH3T3 cells (Wei et al., 2013). Collectively, these studies and our data indicate that IAA triggers oxidative stress and DNA damage in different types of cells. In addition, Collins et al. (2015) and Ding et al. (2019) indicated that DNA damage in mouse oocytes can result in MI arrest that can cause PBE failure. We, therefore, suggest that the IAA-induced DNA damage was one of the causes of MI arrest in the IAA-treated oocytes.

Based on our results, we hypothesized that IAA-induced oxidative stress was the main reason for the oocyte-maturation failure, and supplementation of an antioxidant might rescue the maturation rate in IAA-exposed oocytes. NAC is an antioxidant that can decrease the ROS level and alleviate oxidative stress in cells. Using NAC as an antioxidant can reduce toxicant-induced-oxidative stress and rescue PBE failure in porcine and mouse oocytes (Lai et al., 2015; Li and Zhao, 2019). Our results showed that 200 μM NAC supplementation significantly reduced ROS levels and increased the PBE rate in IAA (5 μM and 10 μM)-exposed oocytes (Fig. 7). This indicates that NAC alleviates the meiotic damage caused by IAA and thereby restores oocyte maturation, which directly confirms that IAA-induced oocyte toxicity is mainly mediated by oxidative stress. Notably, IAA-induced damages in HepG2 cells (Wang et al., 2014) and primary cerebellar granule neurons CGNs (Zhou et al., 2015) were also reduced by antioxidants, which further suggests that oxidative stress is the major mediator by which IAA induces defects in a variety of mammalian cells.

## 5. Conclusion

In conclusion, our data show that IAA exposure impairs meiotic progression and oocyte maturation via disrupting spindle assembly, inducing oxidative stress, and damaging DNA. Moreover, antioxidant NAC supplementation rescues the maturation failure in IAA-exposed oocytes to some extent. This study provides novel information on the toxicological mechanisms underlying the effects of IAA on oocyte maturation and raises concerns of IAA exposure for potential risks of human and animal reproductive health. Although these data suggest that IAA could impair oocyte maturation, further studies are needed to investigate the effects of IAA on oogenesis *in vivo*.

### CRediT author statement

**Xiaofei Jiao:** Conceptualization, Methodology, Validation, Investigation, Formal analysis, Writing-Original Draft. **Andressa Gonsioroski:** Resources, Writing-Review & Editing. **Jodi A Flaws:** Resources, Writing-Review & Editing, Funding acquisition. **Huanyu Qiao:** Conceptualization, Resources, Project administration, Supervision, Writing-Review & Editing, Funding acquisition.

## Acknowledgement

We would like to thank the members of Dr. Flaws’ Laboratory for their help in data collection and kindly sharing reagents. We would like to thank the laboratories of Drs. CheMyong Ko, Michael J Spinella, Sarah Freemantle, and Bo Wang for their technical support. We would also like to thank Qiuman Liang for her help in data analysis. This work was supported by NIH R00 HD082375, NIH R01 GM135549, and NIH R21 ES028963.

## Notes

### Competing Interest Statement

The authors have declared no competing interest.

