## Supplemental Figure 1 for "Iodoacetic acid disrupts mouse oocyte maturation by inducing oxidative stress and spindle abnormalities"

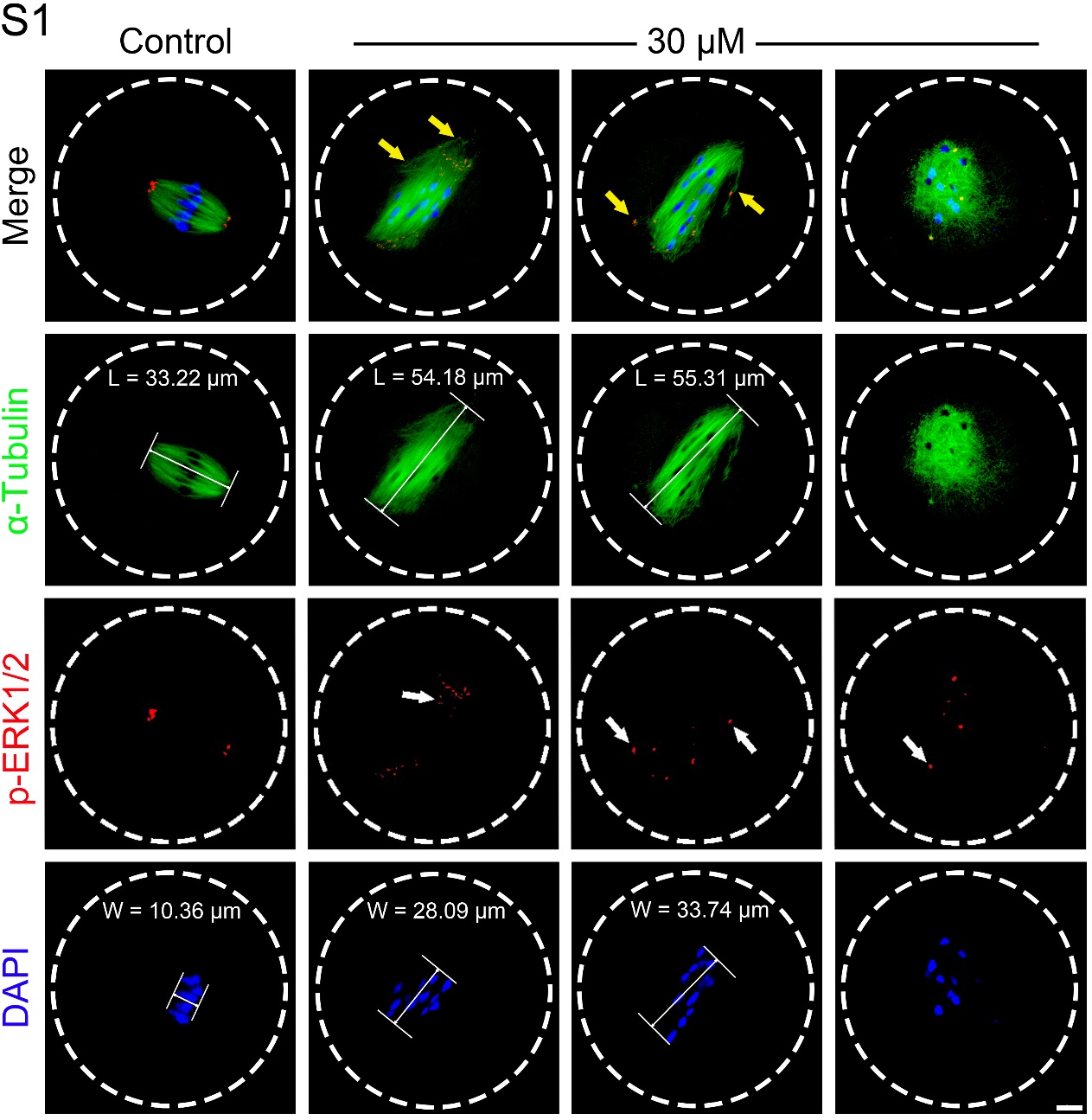

Figure S1

**Figure S1. 30-μM-IAA exposure resulted in severe spindle and chromosome-alignment defects in treated oocytes.** The representative images showed the morphologies of spindles, localization of p-ERK1/2, and chromosome alignment in control and the 30-μM-IAA-treated oocytes. Of note, yellow arrows pointed out the aberrant spindle organization in the IAA-exposed oocytes; the white arrows indicated the mislocalization of p-ERK1/2, microtubule organizing centers (MTOCs) associated proteins that enrich at spindle poles, in IAA-treated oocytes. Spindle length was calculated by the distance from one spindle pole to the other, L = Length. MI plate width was measured by the axis distance between the two lines at the edges of the DNA, W = MI-plate width. α-Tubulin, green; p-ERK1/2, red; chromosomes/DNA, blue. Scale bar = 10 μm.
